# Bill colour outperforms iris colour for sexing in Starlings *Sturnus vulgaris* in breeding condition

**DOI:** 10.64898/2026.09.28.755006

**Authors:** Kristina Pascual, Melissa Bateson

**Affiliations:** Biosciences Institute, Newcastle University, Newcastle upon Tyne, UK

**Author notes:** Correspondence author., Address: Henry Wellcome Building, Framlington Place, Newcastle upon Tyne, NE2 4HH.

## Abstract

Reliable morphological criteria for sexing birds without the need for invasive procedures are scientifically and ethically valuable. Smith *et al* (2005) reported that iris colour assigns the genetic sex of Starlings *Sturnus vulgaris* with 98% accuracy, and bill base colour with 100% accuracy when birds are in breeding condition, but these figures have not been tested independently. We applied their criteria to 31 captive Starlings in breeding condition, using three observers blind to genetic sex, two of them new to Starling sexing. Bill base colour correctly predicted genetic sex for all 29 birds scored, with complete observer agreement. Mean iris score was correct for 27 of 31 (87.1%), fewer than the 94 of 96 reported by Smith et al (2005) for birds in breeding condition (Fisher’s exact test, P = 0.031), although with so few errors in either study this comparison is imprecise. Observer agreement was high, so these errors reflect iris colouration conflicting with genetic sex rather than inconsistent scoring; bill base colour was correct for all four misassigned birds. Body mass and tarsus length classified 77% and 58% of birds correctly, their published thresholds missing in opposite directions. We conclude that bill base colour should be the primary criterion for sexing Starlings in breeding condition; the accuracy of iris colour should not be assumed to be 98%.

## Introduction

Reliably sexing individual birds is a prerequisite for most ringing, behavioural and ecological work, yet many species are monomorphic or only weakly dimorphic based on external characters (Griffiths *et al*. 1998). Genetic sexing resolves this, and PCR-based assays targeting the CHD1 genes are now routine and inexpensive (Griffiths *et al*. 1998, Morinha *et al*. 2012). However, genetic sexing requires tissue, most often blood, and in the UK taking a blood sample from a wild-caught or captive bird is a regulated procedure requiring licensing and justification. Under the refinement principle of the 3Rs (Russell and Burch 1959), a visual criterion that reaches the same answer without the need for an invasive procedure is preferable wherever it is demonstrably reliable. The force of this argument depends on the reliability of the method: a visual criterion adopted on the strength of an accuracy figure that does not hold in the study population exchanges a small welfare cost for an unquantified rate of misclassification with the potential to invalidate scientific conclusions.

The Starling *Sturnus vulgaris* illustrates the problem well. The sexes differ on average in several biometric and plumage characters, but the distributions overlap enough that none is diagnostic in an individual bird; sexing in the hand has therefore relied on the colour of the bill base and the iris. Kessel (1951) visually examined and subsequently dissected 600 North American Starlings with yellow bills. She found that the colour of the base of the lower mandible, pale pink in females and grey-blue in males, yielded what she described as complete accuracy for predicting gonadal sex. She reported the character as usable from November through the breeding season, the period over which the rest of the bill is yellow. She also noted that the pale outer ring of the female iris, although usable throughout the year, was not as reliable, because the ring is sometimes hard to discern in females and a faint ring is occasionally visible in males. Her measurements bear on two commonly taken biometric characters as well: tarsus length did not differ detectably among her four age and sex groups, and of those groups only adult males differed significantly in weight.

Smith *et al* (2005) explored how well these characters predicted genetic sex. Working with 100 wild Starlings caught as juveniles in the South West UK and followed into their first winter, they scored iris colour on a four-point scale describing how light the iris appears relative to the pupil, together with feather length and shape, speckling, body mass and tarsus length, and validated each against genetic sex. Iris colour was the single best predictor, classifying 98% of birds correctly both in August and in February, and in February the colour of the bill base discriminated the sexes with 100% accuracy. Iris colour is accordingly the character that current sexing guidance now puts first. The identification sheets of Blasco-Zumeta & Heinze (2023) give the iris as uniformly dark brown in males and dark with a narrow pale outer circle in females and note that most juveniles can be sexed on iris colour alone, mentioning bill base colour second and only for the breeding season. Guidance for Starlings held in captivity likewise takes the iris scale as the primary criterion citing Smith *et al* (2005) (Bateson and Asher 2010, Bateson 2024).

Two features of Smith *et al*’s (2005) figures deserve attention before they are relied upon. First, they are classification rates from logistic models fitted to the same data that generated them and so describe how well the criteria separate the sample in which they were developed rather than how well they predict sex in an independent sample. Second, they come from one wild population sampled in one year. Iris and bill colouration are continuously varying, state-dependent characters rather than fixed genetic markers, and there is no reason to expect their diagnostic value to be a constant of the species.

Here we report an independent test. We applied the criteria of Smith *et al* (2005), with three observers blind to genetic sex, to a cohort of known-age Starlings hand-reared from wild nests in the North East UK and held in captivity to 11 months of age. We compared the visual assignments with genetic sex. We ask three questions: how accurately do bill and iris colour assign genetic sex in a novel sample; how repeatable are the scores among observers; and where the visual assignment is wrong, does the error lie with the observer or are the characters genuinely discordant in some birds?

## Methods

### Subjects and housing

Subjects were 31 Starlings taken from wild nest boxes in Northumberland, UK (55°15′N, 2°00′W) at four days post-hatching in May 2022 under licence from Natural England (Ref: 2022-56110-SCI-SCI). Birds were hand-reared and subsequently held in indoor aviaries under full-spectrum lighting in mixed-sex groups. They formed the two arms of an unrelated 10-day early-life nutritional manipulation applied from day 6 post-hatching (16 birds benign, 15 birds harsh); after that manipulation ended, all birds were reared and maintained identically (see Pascual 2026 for further details). At the time of visual assessment, the birds were 11 months old and had been held on a short-day photoperiod (10 h light: 14 h dark) for approximately six months. Starlings held on short days become photosensitive and come into breeding condition (Dawson *et al*. 2001) and all birds showed the predominantly yellow bill characteristic of reproductively active Starlings (Witschi and Miller 1938, Wydoski 1964). The 31 birds came from 16 broods: 15 sibling pairs, with one member of each pair allocated to each nutritional condition, and one bird whose sibling was not available.

### Visual assessment

All birds were caught from their home aviary and assessed on a single day (28 April 2023) by three observers (K. Pascual, M. Bateson and E. Maley) working independently and blind to genetic sex, which had not been determined at that point. Each observer scored the iris on the four-point scale of Smith *et al* (2005), which rates how light the iris appears relative to the pupil, from 1 (pale, whitish iris) through 2 (lighter, clear ring) and 3 (dim ring, visible only with careful observation) to 4 (so dark as to be indistinguishable from the pupil); low scores indicate female traits and high scores male traits. Each observer also classified the colour of the base of the lower mandible as pink or grey-blue. Body mass was recorded at the same session to the nearest 0.1 g. Tarsus length had been measured at 123 days post-hatching, to the nearest 0.01 mm using the same maximum tarsus method (Redfern and Clark 2001) as Smith *et al* (2005). Tarsal growth in Starlings is complete by 15 days post-hatching (Allen *et al*. 2022), so this measure reflects adult size.

Following Smith *et al* (2005) we had intended also to score feather length and feather tip shape. Speckling was excluded at the outset because Smith *et al* (2005) found it uninformative. As observers new to the technique, we were unable to score the two plumage characters consistently and abandoned them during the session to reduce bird handling time. Due to an error, two birds received no bill score from any observer, and a third bird was missed by one observer, who recorded neither its iris nor its bill. Bill colour was therefore scored for 29 birds, 28 of them by all three observers, and iris colour for all 31, 30 of them by all three.

### Genetic sexing

Blood was sampled four to eight weeks after visual assessment, on 24 May 2023 (12 birds) and 23 June 2023 (19 birds), under Home Office licence (PPL number: P038AB1D3) and with the approval of the Newcastle University Animal Welfare and Ethical Review Body. Approximately 225 μl of blood was taken by venepuncture into 75 μl capillary tubes, the majority of it for purposes unrelated to this study. A drop from one capillary tube was spotted onto a sampling card and sent to Avigenics (London, UK) for commercial CHD1-based genetic sexing. Body mass was also recorded at this session. Results were returned in July 2023, after all visual scoring was complete. The assay reports the presence or absence of a W-linked CHD1 allele and thus determines genetic sex. We treat its result as the reference standard throughout and describe a visual assignment as correct when it agrees with the determined genetic sex.

### Data analysis

Analyses were carried out in R 4.6.1 (R Core Team 2026). Claude Opus 5.5 was used to assist in writing the final version of the R script to improve clarity and to check that the results were fully reproducible. Following Smith *et al* (2005), we assigned sex from each trait using their rules: a mean iris score of 3 or above indicates a male, a grey-blue bill base indicates a male, a body mass of 78 g or above indicates a male, and a tarsus length of 29.3 mm or above indicates a male. For each trait we report the proportion of birds correctly assigned with a Wilson 95% confidence interval (Brown *et al*. 2001) and we fitted a binomial generalised linear model with genetic sex as the response, reporting each coefficient with a 95% profile-likelihood confidence interval (Venzon and Moolgavkar 1988). We do not report significance tests for these coefficients because with 31 birds the tests available for a logistic coefficient are unreliable. The classification accuracies and areas under the ROC curve, both reported with confidence intervals, provide the statistical inference. Bill colour separated the sexes completely, so no logistic model could be fitted and Fisher’s exact test was used instead.

Discrimination independent of any cut-point is summarised by the area under the ROC curve (Hanley and McNeil 1982), where the AUC equals the probability that a randomly chosen male has a higher score than a randomly chosen female, with ties counted as half (0.5 is no discrimination, 1.0 is complete separation). Confidence intervals for the AUC were obtained by resampling broods rather than individual birds (200,000 replicates), so that the sibling structure of the sample is preserved. From the same resampling we also report, for the characters whose interval includes 0.5, the proportion of replicates in which the AUC was 0.5 or less; this describes how much of the bootstrap distribution lies at or below chance. To ask whether the quantitative characters added anything to iris score, we fitted a model containing iris score, body mass and tarsus length together and compared it with the iris-only model by AIC and by the confidence intervals on the individual coefficients.

Inter-observer reliability was assessed with Kendall’s W for the ordinal iris scores and Fleiss’ kappa for the binary bill scores and for the sex assignments derived from iris score. Since the iris scale has only four levels and ties are pervasive, we report W corrected for ties. To test whether our accuracy for iris colour differed from that reported by Smith *et al* (2005), we compared the two studies’ counts of correct and incorrect assignments in a 2 × 2 table using Fisher’s exact test, taking their February sample as the comparison (that being the occasion on which, as in our study, birds were in breeding condition). They report 98% of 96 birds correctly assigned, which corresponds to 94 of 96. Tests are two-tailed throughout. The data and the analysis script are available at: https://doi.org/10.17605/OSF.IO/ZG56M.

## Results

Genetic sexing identified 17 males and 14 females. Accuracy and model results for each trait are given in Table 1, and the distributions of iris score, body mass and tarsus length by genetic sex are shown in Figure 1.

**Table 1.** Accuracy of each visual trait in assigning the genetic sex of 31 captive Starlings in breeding condition, using the classification rules of Smith *et al* (2005).

| Trait | n | Correct | % correct (95% CI) | b (95% CI) |
| --- | --- | --- | --- | --- |
| Bill base colour | 29 | 29 | 100.0 (88.3-100.0) | - |
| Iris colour (mean score) | 31 | 27 | 87.1 (71.1-94.9) | 2.04 (1.04-3.61) |
| Body mass | 31 | 24 | 77.4 (60.2-88.6) | 0.131 (0.023-0.267) |
| Tarsus length | 31 | 18 | 58.1 (40.8-73.6) | 0.856 (0.019-1.887) |

Notes: Sample sizes differ because two birds received no bill score from any observer and a third was missed by one observer. Wilson confidence intervals are given for % correct. Profile-likelihood confidence intervals are given for the model coefficients (not available for bill colour because it separated the sexes completely and a Fisher's exact test is reported instead, $P < 0.001$ ).

**Figure 1.**
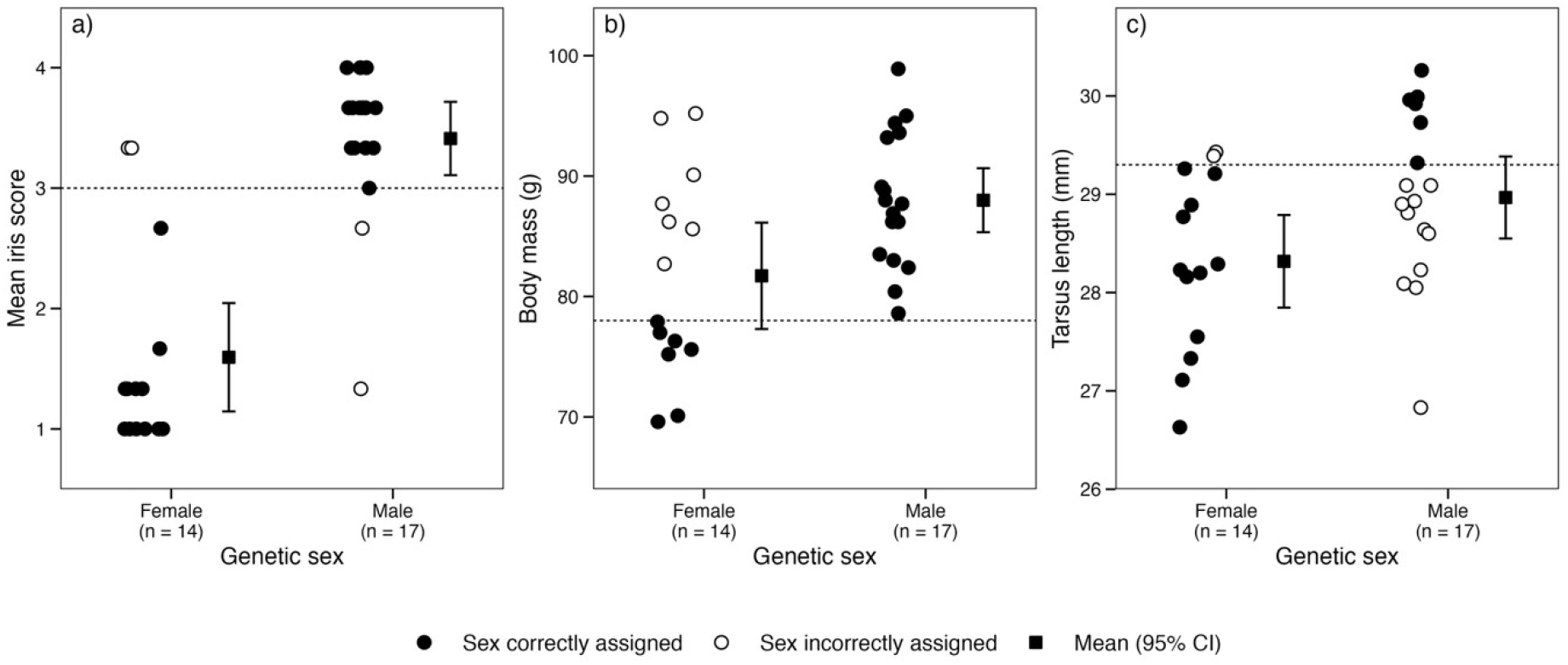
Iris colour, body mass and tarsus length as predictors of genetic sex in 31 captive Starlings in breeding condition: a) mean iris score across three independent observers (two observers for one bird), b) body mass at the time of assessment, and c) adult tarsus length. Dashed lines mark the sex-assignment thresholds of Smith *et al* (2005). Filled circles are birds whose genetic sex that trait assigned correctly and open circles those it assigned incorrectly; squares with error bars are means with 95% confidence intervals. Bill base colour matched genetic sex for every bird scored, including all four birds misassigned by iris score.

### Bill base colour

Of the 29 birds that received a bill score, all 29 were assigned to the correct sex (100%, 95% CI 88.3– 100.0; Fisher’s exact test, P < 0.001). Every bird with a pink bill base was a genetic female and every bird with a grey-blue bill base was a genetic male. The three observers agreed on every bird they all scored (Fleiss’ kappa = 1.00, 28 birds scored by all three), and each observer taken alone was correct on every bird scored.

### Iris colour

Mean iris score was correct for 27 of 31 birds (87.1%, 95% CI 71.1–94.9) and was a strong predictor of genetic sex (b = 2.04, 95% CI 1.04 to 3.61; area under the ROC curve = 0.94, 95% CI 0.87-1.00). As in Smith *et al* (2005), low scores were characteristic of females (mean 1.60, sd 0.86) and high scores of males (mean 3.41, sd 0.64). The four errors were not an artefact of the cut-point: accuracy was 87.1% for every threshold between 2.0 and 3.0, and leave-one-out cross-validation of the fitted model also gave 27 of 31. Individual observers achieved 86.7%, 87.1% and 87.1%.

### Inter-observer reliability

Agreement on iris score was high (Kendall’s W = 0.88 corrected for ties; 30 birds scored by all three observers). All three observers gave an identical score on only nine of 30 birds (30.0%), but no disagreement spanned more than a single scale point, so almost all disagreements were immaterial to the sex assigned: observers agreed on the assigned sex for 28 of 30 birds (93.3%; Fleiss’ kappa = 0.91).

### Misassigned birds

Two genetic males were assigned as female (mean iris scores 1.33 and 2.67) and two genetic females as male (both 3.33). Three of the four were scored identically by all three observers, and for the fourth the disagreement was one point and did not cross the decision threshold; the errors therefore reflect iris colouration that conflicted with genetic sex rather than inconsistent scoring. The four birds came from four different broods and were evenly split between the two early-life treatment groups (two and two; Fisher’s exact test, P = 1.00), although with four errors this sample cannot exclude a modest effect of either factor. In all four cases the bill base colour was correct. Iris and bill gave conflicting assignments for exactly these four birds out of the 29 scored for both, and bill colour was correct on every occasion. Figure 2 shows photos of two misassigned birds.

**Figure 2.**
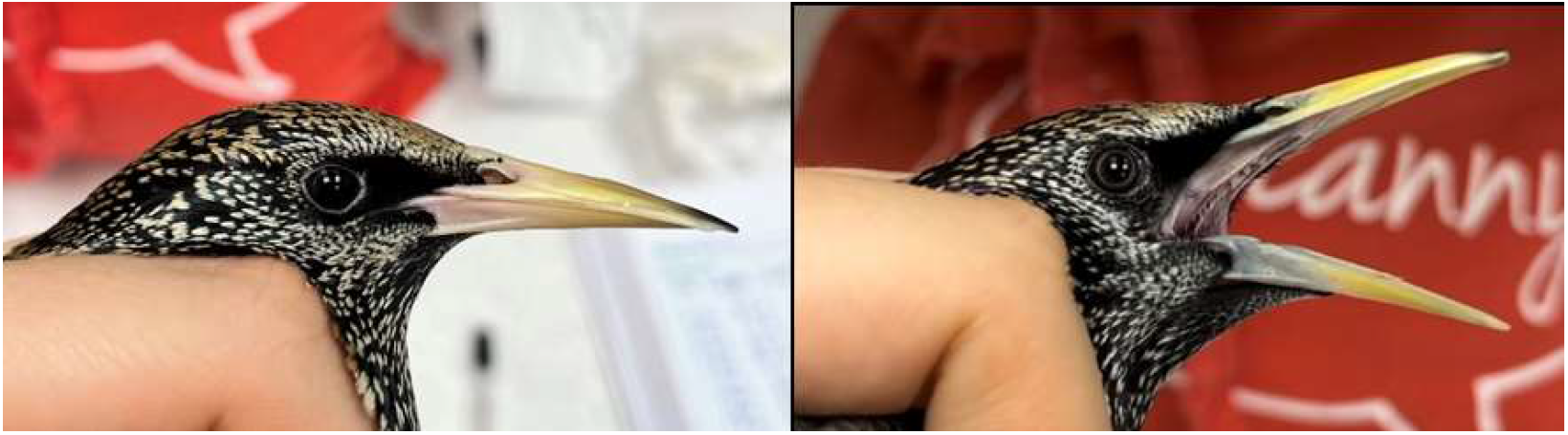
Two of the four birds whose iris colour conflicted with their genetic sex: a) a genetic female with a high mean iris score but a pink bill base, and b) a genetic male with a clearly visible pale ring, and therefore a low mean iris score, but a grey-blue bill base.

### Body mass

Applying the 78 g threshold classified 24 of 31 birds correctly (77.4%, 95% CI 60.2–88.6). Mass predicted sex (b = 0.131, 95% CI 0.023 to 0.267) but discriminated poorly (area under the ROC curve = 0.72, 95% CI 0.53-0.89), and the distributions overlapped substantially (females mean 81.7 g, sd 8.4; males mean 88.0 g, sd 5.6; Figure 1b). Seven of the 14 females exceeded the 78 g threshold and no male fell below it, so all seven errors were females classified as male. The threshold from Smith *et al* (2005) is therefore too low for these birds. Since body mass in Starlings varies within and between individuals (e.g. Bateson and Nolan 2022), no fixed threshold should be expected to transfer between samples.

### Tarsus length

Applying the 29.3 mm threshold classified 18 of 31 birds correctly (58.1%, 95% CI 40.8-73.6). Tarsus length predicted sex weakly (b = 0.856, 95% CI 0.019 to 1.887) and discriminated poorly (area under the ROC curve = 0.68, 95% CI 0.48-0.87; Figure 1c). The threshold is above the mean tarsus length of our males (females mean 28.32 mm, sd 0.90; males mean 28.97 mm, sd 0.88), and 11 of the 13 errors were males classified as female. Smith *et al* (2005) do not report tarsus means, so we cannot say directly that our birds were smaller than theirs; what the data show is that a threshold which discriminated 65-67% of their birds is not where it needs to be in this sample. Re-deriving the cut-point within our own sample would raise accuracy to 67.7% (at 28.6 mm). The confidence interval on the AUC includes 0.5, so at this sample size tarsus length cannot be firmly distinguished from a character carrying no information about sex at all: 4% of the bootstrap replicates gave an AUC of 0.5 or less. Kessel (1951) likewise found no detectable difference in tarsus length among her age and sex groups.

### The characters considered together

Iris score, body mass and tarsus length taken together predicted sex no better than iris score alone. Adding mass and tarsus length to the iris-only model raised AIC from 24.3 to 28.1, and the confidence intervals on both coefficients span zero once iris score is in the model (mass: −0.298 to 0.182; tarsus −1.395 to 2.079), while that on iris score does not (0.930 to 4.227).

### Comparison with Smith *et al* (2005)

Our accuracy for iris colour, 27 of 31 correct, was lower than the 94 of 96 achieved for birds in breeding condition by Smith *et al* (2005) (Fisher’s exact test, P = 0.031); the same test against their August sample of 98 of 100 gives P = 0.028. The corresponding confidence interval for the odds ratio is very wide and includes one (odds ratio 6.83, 95% CI 0.92–79.2), reflecting how few errors either study recorded, so the difference should be regarded as suggestive only. Testing our proportion against a fixed expectation of 98% gives the same conclusion more strongly (exact binomial test, P = 0.003), but that test treats the published figure as known without error and we prefer the more conservative comparison. Bill colour, at 29 of 29 against 96 of 96, is indistinguishable between the two studies.

## Discussion

Applying the criteria of Smith *et al* (2005) to an independent cohort reproduced one of their two main results and qualified the other. Bill base colour was perfectly diagnostic, as it was for them (and as it was for Kessel (1951) based on gonadal sex). It was also, in our data, the easiest character to use: three observers, two of them new to Starling sexing, agreed on every bird, with no training beyond the published description. For Starlings in breeding condition, we therefore endorse the existing recommendation without reservation and note that it is the character least likely to be affected by lighting, observer experience or the condition of the bird.

Iris colour behaved differently. It remained a strong predictor of sex, but it assigned 87% of birds correctly rather than 98%, and the shortfall does not appear to be a consequence of our scoring. Agreement among observers was high, no disagreement spanned more than one point on the scale, accuracy was identical for every cut-point we could reasonably have chosen, and each observer independently achieved the same accuracy as the group, the one experienced observer no better than the two new to the technique. Three of the four misassigned birds were scored identically by all three observers. The most parsimonious interpretation is that these four birds had iris colouration that genuinely did not match their genetic sex. That is not a new phenomenon: Kessel (1951) noted that the pale ring is sometimes indiscernible in gonadal females and sometimes faintly present in gonadal males.

Our discussion thus far assumes the genetic sexing result is correct, which deserves examination. The assay reports the presence or absence of a W-linked CHD1 allele, and its commonest failure is that the W band does not amplify, turning a genetic female (ZW) into an apparent male (ZZ). Two of our four discordant birds are genetic males whose iris scores indicated female, which is exactly what such a failure would produce. However, bill base colour argues against that explanation: both birds were scored grey-blue by all three observers, and the two discordant genetic females were both scored pink, so on every bird where iris colour disagreed with the genetic sex an independent phenotypic character (bill base colour) supported the genetic sex. The same holds across the dataset: bill colour matched the genetic result for all 29 birds scored, which would be unlikely had samples been mislabelled or assays failed at any appreciable rate. Bill base colouration is under gonadal hormonal control, so its agreement suggests that the causal link from genotype through gonad to phenotype is intact in these birds, and that in these four it is specifically iris colour that departs from it.

Why our rate of such birds should be higher than that of Smith *et al* (2005) we cannot establish from these data, and we are wary of over-interpreting a difference resting on four errors against two. Several explanations are compatible with what we observed. The two samples differ in population, in year and in rearing environment: our birds were taken from the nest at four days old and reared by hand in captivity, whereas theirs were wild-caught juveniles. They also differ in age and photoperiodic history, our birds being 11 months old and brought into breeding condition on short days rather than in their first winter under natural photoperiod. A further possibility has nothing to do with Starlings: Smith *et al* (2005) report the classification rate of a model fitted to their own data, which is an optimistic estimate of how a criterion will perform out of sample, whereas we applied a fixed rule to a new sample. Our own figure is an out-of-sample estimate, since the rule was fixed before these birds were scored, and leave-one-out cross-validation of a model fitted to our own data returned the same 27 of 31, so nothing in our estimate is inflated by fitting. Some part of the gap between 87% and 98% is likely to be the ordinary difference between those two quantities. We found no evidence that misassignment was associated with early-life nutritional treatment or with nest of origin, but with four errors the power to detect such associations is negligible.

Neither body mass nor tarsus length was of much use on its own for predicting genetic sex, and neither added anything to iris score. What is useful is the pattern of errors. The 78 g mass threshold is too low for our birds, so every mass error was a female classified as male; the 29.3 mm tarsus threshold is too high, so almost every tarsus error was a male classified as female. Neither cut-off is locating the boundary between the sexes, and because they miss in opposite directions in the same 31 birds this cannot simply be due to our birds being larger or smaller overall. Neither result is surprising: Kessel (1951) found that tarsus length did not separate her age and sex groups at all, and body weight separated only adult males.

The practical implications for ringers are straightforward. Where birds are in breeding condition, bill base colour should be the primary criterion: it was correct for every bird in this study and for every bird in the February sample of Smith *et al* (2005), and it was correct for all four birds whose iris colour was misleading. The time window during which bill base colour is useful is also wider than the breeding season itself, since Kessel (1951) found it worked for predicting gonadal sex from November onwards, once the bill has begun to yellow. Outside that window, when the bill has darkened after moult and the character is unavailable, iris colour remains the best visual criterion, but users should plan on the basis of an accuracy in the high eighties, rather than 98%, and consider the consequences for their design. Where a few percent of misassigned individuals would not matter, visual sexing is an appropriate refinement that avoids a licensed procedure; where conclusions depend on a sex difference, particularly with modest sample sizes, an error rate of one bird in eight is not negligible and genetic confirmation is warranted. Where both characters are available and they disagree, our data, though only four birds out of 29, favour the bill without exception. For birds in breeding condition this reverses the order of priority in current guidance, which puts iris colour first and treats bill base colour as a secondary character available only in the breeding season (Blasco-Zumeta and Heinze 2023). Our four iris errors also qualify the caution that accompanies that guidance, namely that both sexes may show a uniformly dark iris but only females the pale ring. The two genetic females we assigned as male had dark eyes in which any ring was at best dim, which is the ambiguity the caution anticipates; but the two genetic males we assigned as female showed the ring that the same guidance treats as diagnostic of females, so a visible ring is not by itself sufficient evidence that a bird is female.

A practical observation on the remaining characters is also worth recording. We abandoned the two plumage characters during scoring because we could not apply them consistently. Smith *et al* (2005) report that feather length classified 93–94% of birds correctly, a useful figure but one obtained by more experienced observers. The transferability of a visual criterion to a novice observer is a separate property from its accuracy in expert hands. Bill colour required no such expertise, which is a further argument for its primacy.

Four limitations should temper these conclusions. The first is sample size: 31 birds gives a confidence interval on our iris accuracy running from 71% to 95%, consistent with a true accuracy well below the previously published figure. The second is that these 31 birds are not 31 independent samples. They came from 16 broods, 15 of them contributing a pair of siblings, and if iris colouration has any heritable or shared-environment component then our effective sample size is smaller than 31 and the confidence intervals reported here are correspondingly too narrow. The four misassigned birds did at least come from four different broods, so the shortfall in accuracy is not the product of one atypical family. The third is that our birds were captive-reared and we cannot say how our estimate transfers to wild birds. The fourth applies to the biometric characters alone: half of these birds underwent an early-life nutritional manipulation that affected growth and body mass, reported in full elsewhere, so the sample may not be representative of an unmanipulated cohort. We therefore present the mass and tarsus results as a demonstration that imported thresholds need not transfer, rather than as an estimate of how those thresholds would perform in wild birds. The colour characters are not affected in the same way: iris errors were divided evenly between the two treatment groups. What these results establish is not that the published accuracy of iris colour is wrong, but that it is not a property of the species that can be assumed: it is an estimate from one sample, and at least one other sample gives a materially lower value. That distinction matters for any character whose purpose is to justify replacing a regulated procedure, and it applies beyond just Starlings, since sexing and ageing criteria across many species rest on single validation studies whose accuracies are quoted thereafter as though they were constants. Ringers and laboratories that hold paired visual and genetic or gonadal sex determinations should report them, since only the accumulation of such estimates will establish the accuracy of a visual criterion.

## Acknowledgements

We thank Ella Maley for her help with data collection, and Michelle Waddle for assistance with bird husbandry. The project was part-funded by a grant from the Wild Animal Initiative.

## Data availability

The data and R code supporting this study are openly available in the Open Science Framework at:

https://doi.org/10.17605/OSF.IO/ZG56M

